# Cyclic immunofluorescence platform using photocleavable linkers for direct antibody labeling enables cancer phenotyping

**DOI:** 10.64898/2026.07.19.735577

**Authors:** Amanda Zucker, Cam Nguyen, Frederik Brøndsted, Jocelyn A. Jones, Gauri S. Malankar, Taelor J. Ekstrom, Cody Rounds, Divya Ravi, Shaun M Goodyear, Adel Kardosh, Melissa H. Wong, Lei G. Wang, Summer L. Gibbs

## Abstract

Advances in spatial proteomics through the development of multiplexed immunostaining platforms have facilitated analyses with increasing cellular and molecular granularity. However, currently available approaches are limited by harsh conditions for signal removal, restricting the number of antigens that can be probed in a single specimen without significant alterations to sample quality and structure. Here we present an approach for direct labeling of primary antibodies with fluorophores using a photocleavable linker (PCL) with a polyethylene glycol spacer (PEG) to enable cyclic immunofluorescence (cyCIF) with gentle signal removal conditions. Our innovative approach uses directly labeled primary antibodies to enhance staining specificity and cyclic immunostaining efficiency, while minimizing nonspecific background signal. Additionally, through integration of the PCL, this approach facilitates gentle cleavage of antibody conjugated fluorophore, preserving sample integrity over multiple rounds of staining. Direct PEG-PCL antibody labeling will promote greater multiplexing by minimizing specimen damage and allow for quantitative analyses of cyCIF spatial data. We demonstrate that cyCIF with PEG-PCL conjugated antibodies can be applied across a variety of cancer subtypes to identify and characterize rare neoplastic cell populations in both tumor tissue and fragile peripheral blood specimens.

## Introduction

Novel scientific discoveries and therapeutic innovations have reshaped oncology, yet cancer remains among the leading causes of death worldwide. By 2050, the number of new cancer diagnoses is expected to reach 35 million annually^1, 2^. Diagnostics rely on a combination of clinical imaging, tissue biopsy, and surgery to localize tumors and identify specific cancer subtypes facilitating personalized therapy. One of the barriers to effective cancer treatment is the lack of phenotypic analyses, providing insight into structural organization and cellular signaling networks to stratify disease heterogeneity for optimal therapy. Quantification of cancer phenotypes can highlight potential therapeutic vulnerabilities, mechanisms of resistance, and enable prognostic assessment of disease progression. Integrating phenotypic analyses in diagnostic testing and disease monitoring is a promising strategy to improve treatment decisions throughout the clinical course and ultimately patient outcomes.

A variety of molecular imaging tools are under development to assess phenotypes and protein expression of heterogeneous cell populations across a myriad of cancer types. Traditional immunofluorescence (IF) and immunohistochemistry (IHC) are powerful tools for detection and visualization of disease biomarkers. However, both traditional IF and IHC are limited by the number of biomarkers that can be analyzed on a single sample. Therefore, multiplexed immunostaining techniques have been developed facilitating spatial proteomics. Highly multiplexed imaging technologies, including mass-spectrometry imaging (MSI), multiplexed immunohistochemistry (mIHC) and cyclic immunofluorescence (cyCIF), facilitate analyses of spatial biology at the single-cell level to understand disease mechanisms across cancer types. Implementation of MSI technologies have been increasingly common in research to uncover disease biology due to their high sensitivity and utility for multiplexing. MSI techniques, including cytometry by time-of-flight (CyTOF) and multiplexed ion beam imaging (MIBI), utilize antibodies labeled with rare earth metals with demonstrated simultaneous detection of >40 protein targets^3-5^. While this enhances multiplexing, cellular identification^4, 6, 7^, and single-cell image analyses, multiplexed immunostaining using MSI technologies are limited by instrument accessibility for clinical use, challenging antibody labeling requirements, and reduced spatial resolution due to the scanning laser spot size^5, 8-10^. In contrast to MSI, mIHC and cyCIF utilize standard microscopy instrumentation. However, mIHC remains limited in the number of markers detectable on the single sample and requires harsh signal removal techniques such as antibody stripping.

To enhance IF performance and the number of biomarkers that can be assessed in a single sample, multiplexed immunofluorescence imaging technologies have been developed to utilize cycles of fluorescent tagging, imaging, and signal removal through chemical bleaching or dissociation of affinity tags^11-16^. However, many cyCIF techniques remain limited by the harsh conditions required for signal removal (e.g., fluorophore photobleaching) which compromise sample integrity and complicate image co-registration as cycle number increases^17^. To overcome these limitations, we previously developed a flexible cyCIF methodology using DNA barcoded antibodies (i.e., oligonucleotide labeled antibodies) using single stranded DNA, where antigen detection was completed by in situ hybridization of the fluorophore labeled complementary oligonucleotide strand. For gentle signal remove, a photocleavable linker (PCL) was incorporated into the complementary fluorophore-labeled single stranded oligonucleotide, facilitating signal removal through ultraviolet (UV) light exposure^18-20^. Our DNA barcoded antibody platform (Ab-oligo cyCIF) minimized total staining time with the addition of all primary oligonucleotide conjugated antibodies (Ab-oligos) in a single round, followed only by addition of imaging strand between rounds of imaging and fluorophore cleaving. However, cyCIF with Ab-oligos can result in non-specific background due to unhybridized oligonucleotides^19^. Specifically, complementary fluorophore labeled oligonucleotides used to detect Ab-oligo conjugates are often negatively charged and thus attracted to the positively charged nucleus, resulting in nonspecific nuclear signal. This nonspecific nuclear signal is amplified by the net negative charges of the fluorescent dye labels, creating challenges in quantification of nuclear protein targets. Nonspecific nuclear signal can be minimized through specific staining methodologies, but overall, the Ab-oligo cyCIF method performs best for membrane and cytoplasmic biomarkers with high expression, while biomarkers with relatively low expression are more difficult to detect^19^.

In this study, we demonstrate that PCL-fluorophore conjugation can be applied to primary antibodies for cyCIF applications with direct antibody staining, overcoming the limitations faced with Ab-oligo cyCIF. Direct antibody labeling reduces overall staining time, permits spatial visualization of endogenous protein expression, enhances staining specificity, and facilitates protein expression quantification for each antibody target, while preserving sample integrity. With direct antibody-fluorophore conjugation no indirect detection method is required, removing previous difficulties with nonspecific signal from unhybridized oligonucleotide labeling strands. Additionally, directly labeled antibodies generally emit brighter fluorescence signal than oligonucleotide labeled antibody conjugates, making directly conjugated antibodies better suited for detection of low abundance biomarkers^21^. Our novel cyCIF technology incorporates PCL-fluorophore conjugates with direct primary antibody labeling, where we optimized polyethylene glycol (PEG) spacer length to minimize photoinduced electron transfer (PET) and the resulting fluorophore quenching due to direct interaction of the PCL and fluorophore. Our direct conjugation approach using PEG-PCL conjugated fluorophores maintains the benefits of UV photocleavage of fluorophores from our Ab-oligo cyCIF methodology with enhanced target specificity and antigen labeling brightness. Here, we validate that cyCIF with PEG-PCL-fluorophore conjugated antibodies can be successfully implemented with primary tumors across cancers and patient peripheral blood samples to identify, phenotype, and classify single cells by protein expression. We performed proof of concept staining using lung squamous cell carcinoma (LUSCC), head and neck squamous cell carcinoma (HNSCC), pancreatic ductal adenocarcinoma (PDAC) and pancreatic adenosquamous carcinoma (PASC) to validate that PEG-PCL mediated cyCIF facilitates detailed annotation of the tumor spatial biology and cellular phenotypes. PEG-PCL cyCIF analyses of tumors across three organ sites demonstrated our ability to identify histologically distinct regions of tumor, immune infiltrate, and cell signaling pathways. We applied the PEG-PCL cyCIF platform to identify and define characteristics of rare cell populations, including hybrid cells. Hybrid cells are a unique cell population formed as a result of fusion between a tumor epithelial cell and an immune cell which are termed circulating hybrid cells (CHCs) when disseminated into peripheral blood^22, 23^. Using PEG-PCL cyCIF, we demonstrated that liquid biopsy using blood draws is feasible, where the relative number of each hybrid cell type in the blood reflected the tumor-based classifications. Thus, cyCIF analysis of patient blood to identify and phenotype CHCs provides a platform to infer characteristics of the primary tumor that drive oncogenesis and metastatic progression to inform clinical decisions.

## Methods

### Human specimens

All human tissue specimens were collected and analyzed under approved protocols in accordance with the ethical requirements and regulations of the institutional review board at Oregon Health & Science University (OHSU). 5 µm formalin-fixed paraffin-embedded (FFPE) tissue sections from pancreatic ductal adenocarcinoma (PDAC), pancreatic adenosquamous carcinoma (PASC), head and neck squamous cell carcinoma (HNSCC), and lung squamous cell carcinoma (LUSCC) were obtained from the OHSU Knight Biolibrary.

Peripheral blood was collected in sodium heparin tubes (BD Vacutainer #367874). Peripheral blood mononuclear cells (PBMCs) were isolated from a minimum of 10 mL of peripheral blood by Ficoll-Paque density gradient^24^, resuspended in fluorescence activated cell sorting (FACS) buffer (deionized PBS, 2% FBS, 1mM EDTA) and adhered at a density of 300,000 PBMCs per well across 3 wells on a poly-D-lysine (0.56 mg/mL; EMD Millipore Corp, USA) coated Superfrost slide (Thermo Fisher Scientific, Waltham, MA). Adhered PBMCs were incubated at 37 °C for 15 minutes, fixed with 4% paraformaldehyde (PFA) and permeabilized with CSK (cytoskeleton) buffer containing 0.5% Triton X-100 (300 mM Sucrose, 100mM NaCl, 30mM MgCl_2_, 10mM PIPES), refixed with 4% PFA, and washed with 2x SSC (sodium chloride-sodium citrate) buffer (G Biosciences #R019). Fixed PBMCs were dehydrated in graded ethanol baths (70%, 95%, 100%).

### General

All reagents were purchased from Sigma Aldrich (St. Louis, MO), Fisher Scientific (Waltham, MA), Ambeed (Buffalo Grove, IL), Chemscene (Monmouth Junction, NJ), and Tokyo Chemical Industry (Portland, OR) and used without additional purification unless otherwise indicated. Analytical thin layer chromatograph (TLC) was performed on Millipore ready-to-use plates with silica gel 60 (F254, 32-63 mm). Purification was performed on a Biotage Isolera Flash System using pre-packaged silica gel cartridges or on a reverse phase preparative HPLC (Agilent 1250 Infinity HPLC).

Mass-to-charge ratio (m/z) and purity of all synthesized compounds were characterized on a tandem Agilent 6244 time-of-flight LCMS with diode array detector VL+. Exact and observed masses are reported (Table 1). Purity of each derivative was quantified via area under the curve (AUC) analysis of the absorbance at 254 nm and m/z ratio in positive ion mode. Sample (5 µL) was injected onto a C18 column (Poroshell 120, 2.1 × 50 mm, 2.7 micron) and eluted with a solvent system of A (H_2_O, 0.1% formic acid) and B (Acetonitrile, 0.1% formic acid) at 0.4 mL/min from A/B = 95/5 to 5/95 over 6 minutes and maintained at A/B = 5/95 for an additional 2 minutes. Ions were detected in positive ion mode by setting the capillary voltage at 4kV and gas temperature at 350 °C.

**Table 1.** Liquid chromatography-mass spectrometry (LCMS) data highlighting structure names, chemical formulas, and exact and observed masses for each compound.

| Name | Chemical Formula | Exact Mass | Observed Mass |
| --- | --- | --- | --- |
| PEG10-PCL | C<sub>36</sub>H<sub>63</sub>N<sub>2</sub>O<sub>18</sub><sup>+</sup> [M+H]<sup>+</sup> | 811.4 | 811.4 |
| PEG2-PCL | C<sub>20</sub>H<sub>31</sub>N<sub>2</sub>O<sub>10</sub><sup>+</sup> [M+H]<sup>+</sup> | 459.2 | 459.2 |
| AF488-PEG10-PCL-COOH | C<sub>57</sub>H<sub>73</sub>N<sub>4</sub>O<sub>28</sub>S<sub>2</sub><sup>-</sup> [M]<sup>-</sup>-1 | 1325.4 | 1325.3 |
| AF488-PEG10-PCL-NHS | C<sub>61</sub>H<sub>75</sub>N<sub>5</sub>O<sub>30</sub>S<sub>2</sub> [M-3H]<sup>-</sup>-2 | 710.7 | 710.6 |
| AF488-PEG2-PCL-COOH | C<sub>41</sub>H<sub>43</sub>N<sub>4</sub>O<sub>20</sub>S<sub>2</sub> [M]<sup>+</sup> | 975.2 | 975.2 |
| AF488-PEG2-PCL-NHS | C<sub>45</sub>H<sub>46</sub>N<sub>5</sub>O<sub>22</sub>S<sub>2</sub><sup>+</sup> [M]<sup>+</sup> | 1072.2 | 1072.2 |
| AF555-PEG10-PCL-COOH | C<sub>70</sub>H<sub>101</sub>N<sub>4</sub>O<sub>31</sub>S<sub>4</sub> [M-4H]<sup>-</sup>-3 | 540.5/540.8 | 540.8 |
| AF555-PEG10-PCL-NHS | C<sub>74</sub>H<sub>104</sub>N<sub>5</sub>O<sub>33</sub>S<sub>4</sub> [M-4H]<sup>-</sup>-3 | 572.9/573.2 | 573.2 |
| AF647-PEG10-PCL-COOH | C<sub>72</sub>H<sub>104</sub>N<sub>4</sub>O<sub>31</sub>S<sub>4</sub> [M-3H]<sup>-</sup>-2 | 824.3 | 824.7 |
| AF647-PEG10-PCL-NHS | C<sub>76</sub>H<sub>107</sub>N<sub>5</sub>O<sub>33</sub>S<sub>4</sub> [M-3H]<sup>-</sup>-2 | 872.8/873.3 | 873.3 |
| AF750-NHS | C<sub>42</sub>H<sub>50</sub>N<sub>3</sub>O<sub>16</sub>S<sub>4</sub> [M-2H]<sup>-</sup>-1 | 980.2 | 980.2 |
| AF750-PEG10-PCL-COOH | C<sub>74</sub>H<sub>106</sub>N<sub>4</sub>O<sub>31</sub>S<sub>4</sub> [M-3H]<sup>-</sup>-2 | 837.3/837.8 | 837.8 |
| AF750-PEG10-PCL-NHS | C<sub>78</sub>H<sub>109</sub>N<sub>5</sub>O<sub>33</sub>S<sub>4</sub> [M-3H]<sup>-</sup>-2 | 885.8/886.3 | 886.4 |

Stock solutions of PEG-PCL conjugated fluorophore constructs (PEG-PCL-FL) were prepared in anhydrous dimethyl sulfoxide (DMSO) at a concentration of 1-10 mM. Reference corrected absorption and fluorescence emission spectra were collected using a SpectraMax M5 spectrometer (Molecular Devices, Sunnyvale, CA). All reported data represent the average of at least n=3 replicate measurements.

### Molar extinction coefficient and quantum yield

Molar extinction coefficients were calculated by diluting PEG-PCL-FL constructs to 1-5 µM in phosphate buffered saline (PBS, pH 7.4, 1% DMSO) and measuring reference corrected absorbance spectra. Absorbance measurements were collected in triplicate, and the average value was reported (Supplementary Table 1). The quantum yields of all fluorophores were measured using samples with absorbance values ranging from 0.07-0.09. The relative quantum yields of the PEG-PCL-FL constructs were calculated using their corresponding parent fluorophores as reference standards^25^. Relative quantum yields were measured in triplicate, and the average was reported (Supplementary Table 1).

### Antibody conjugation

Antibodies were purchased from AbCam (Cambridge, UK), Cell Signaling Technology (Danvers, MA), Biocare Medical (Pacheco, CA), Santa Cruz Biotechnology (Dallas, TX), and Jackson ImmunoResearch (West Grove, PA) to the following targets: ΔNp63 (p40), CD45, pan-cytokeratin (panCK), cytokeratin 5 (CK5), cytokeratin 8 (CK8), epithelial cell adhesion molecule (EpCAM), E-cadherin (ECAD), epidermal growth factor receptor (EGFR), phosphorylated EGFR (pEGFR), phosphorylated mitogen-activated protein kinase (pMEK), and donkey anti-mouse (dMs) secondary. Primary antibodies and the donkey anti-mouse (dMs) secondary antibody (Table 2) were conjugated to fluorophores containing a photocleavable linker and a polyethylene glycol spacer (PEG-PCL) via standard *N-*hydroxysuccinimide (NHS) ester reactions using a molar excess of 5-15 fluorophore to antibody molecules. The reaction was carried out in slightly basic (∼pH 8) 1X PBS for 3 h shaken at room temperature. Amicon 0.5 mL 10 kDa spin filters (Merck Millipore, Burlington, MA) were used for purification. Conjugation ratios were calculated using the maximum absorbance at 280 nm for the antibody and the maximum absorbance of each Alexa Fluor (AF) dye. All PEG-PCL-FL conjugates were validated for functionality in tissues with known biomarker positivity prior to implementation in cyCIF experiments.

**Table 2.**
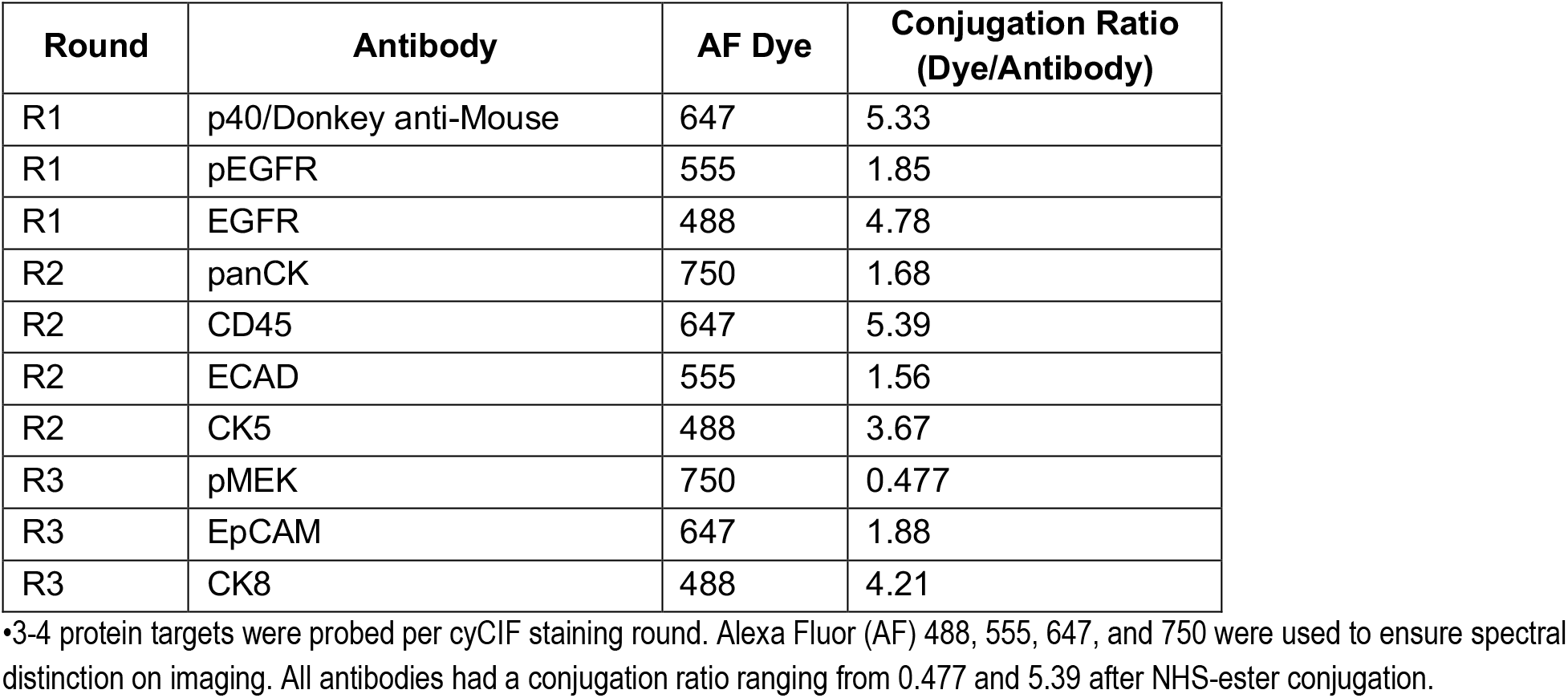
Cyclic immunofluorescence (cyCIF) staining paradigm.

| Round | Antibody | AF Dye | Conjugation Ratio<br>(Dye/Antibody) |
| --- | --- | --- | --- |
| R1 | p40/Donkey anti-Mouse | 647 | 5.33 |
| R1 | pEGFR | 555 | 1.85 |
| R1 | EGFR | 488 | 4.78 |
| R2 | panCK | 750 | 1.68 |
| R2 | CD45 | 647 | 5.39 |
| R2 | ECAD | 555 | 1.56 |
| R2 | CK5 | 488 | 3.67 |
| R3 | pMEK | 750 | 0.477 |
| R3 | EpCAM | 647 | 1.88 |
| R3 | CK8 | 488 | 4.21 |

•3-4 protein targets were probed per cyCIF staining round. Alexa Fluor (AF) 488, 555, 647, and 750 were used to ensure spectral distinction on imaging. All antibodies had a conjugation ratio ranging from 0.477 and 5.39 after NHS-ester conjugation.

### Cyclic immunofluorescence staining

FFPE tumor sections for cyCIF staining were baked at 65 °C for 1.5 hours prior to deparaffinization with xylene washes and rehydration with graded ethanol baths (100%, 95%, 70%, 50%). Tissue underwent antigen retrieval by 30-minute incubation in citrate buffer at 100 °C, a rinse in diH_2_O at 100 °C, 10-minute incubation in Tris-HCl buffer at 100 °C and then cooled to 25 °C with diH_2_O. The PBMC specimen was washed with PBS 3x (5 minutes/wash). Prior to staining with our cyCIF workflow (Figure 1A), tissue and PBMC specimens were counterstained with DAPI and cover-slipped with Fluoromount G to capture background autofluorescence signal as round 0 (R0). Prior to antibody staining, specimens were incubated with blocking buffer containing 1% bovine serum albumin in PBS at 25 °C for 30 minutes. Three to four protein targets were probed during each round of staining. The p40 primary antibody was diluted in blocking buffer (1:75) and was applied to each specimen for overnight incubation at 4 °C in a humidified chamber protected from light. The dMs-PEG-PCL-FL conjugate was diluted to 0.35 µM in blocking buffer and incubated in the dark at 25 °C for 1 hour. The remaining PEG-PCL-FL antibody conjugates (Table 2) were diluted in blocking buffer to a concentration of 15 µg/mL and applied to each specimen, protected from light in a humidified chamber for overnight incubation at 4 °C. Specimens were counterstained with DAPI for 5 minutes at room temperature (RT). Three 5-minute PBS washes were performed between rounds of incubation. Lastly, specimens were cover-slipped with Fluoromount G and imaged using a Zeiss AxioScan Z1 microscope (Carl Zeiss AG, Oberkochen, Germany). An appropriate focus range was determined for each sample individually and exposure times were determined for each antibody. Following each round of imaging, fluorescent signal was cleaved by exposing specimens to UV light for 15 minutes, slides were soaked in 1x PBS for coverslip removal and washed 3x (5 minutes/wash) to remove the cleaved fluorophores before beginning the next round of staining.

**Figure 1.**
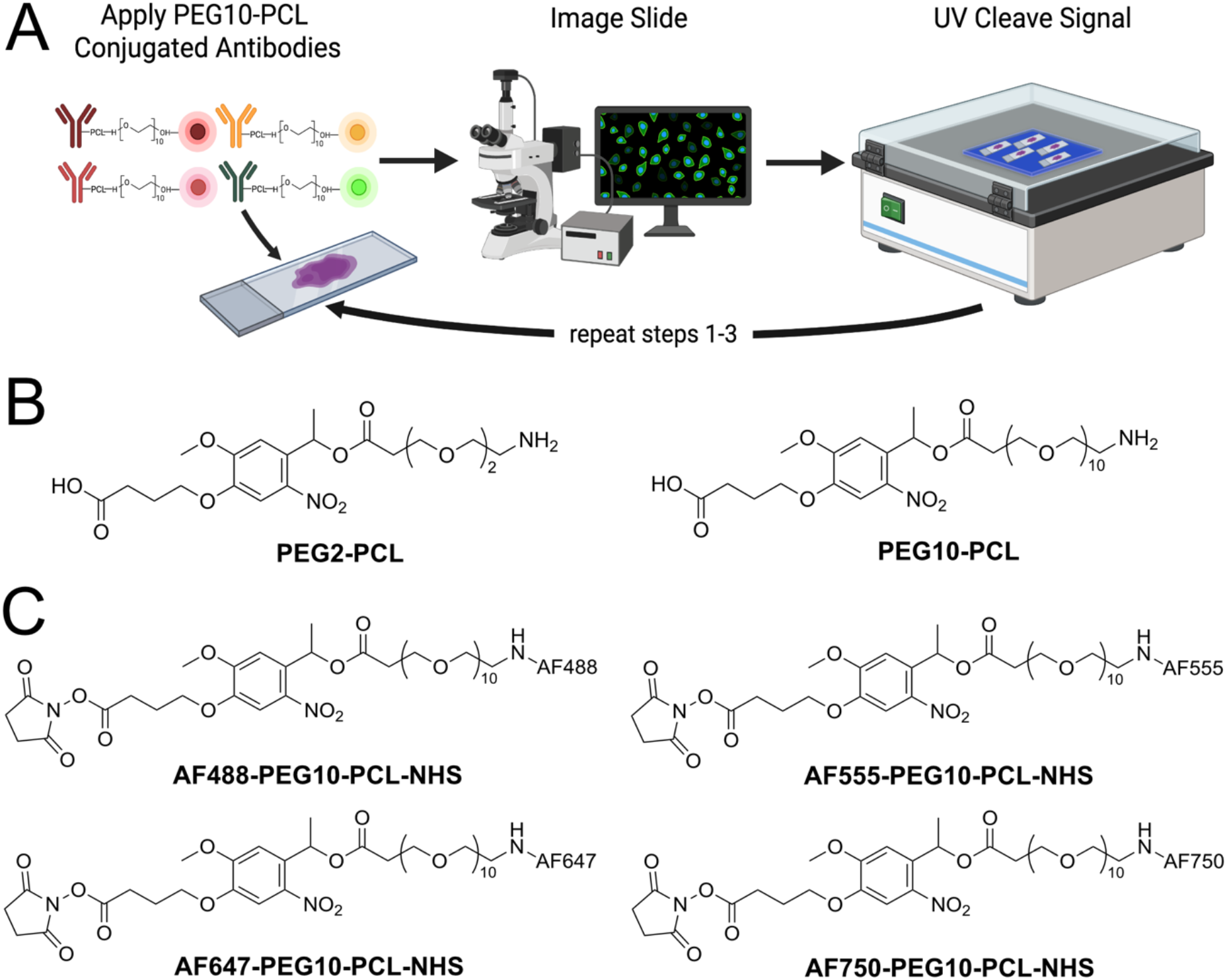
A) Cyclic immunofluorescence (cyCIF) workflow in which up to four spectrally distinct PEG-PCL-FL conjugated antibodies are applied to tissue and blood specimens, imaged, and cleaved with ultraviolet (UV) light. After UV cleavage of fluorophores, coverslips are removed and fluorophores are washed away to prepare for application of the next round of PEG-PCL-FL conjugated antibodies. Figure 1A was created in BioRender. B) Chemical structures of PEG2 and PEG10 spacers used for conjugation to Alexa Fluor (AF) dyes to identify the linker length that results in minimal quenching. C) Chemical structures of PEG10-PCL conjugation to AF488, AF555, AF647, AF750 used for cyCIF applications.

### Image registration and analysis

ASHLAR feature-based registration was implemented to align cyCIF images from each round of staining^26^. Background subtraction was performed on each registered specimen to normalize excitation intensities and exposure times between imaging rounds. Background subtracted images were uploaded to QuPath (version 0.6.0) for downstream hybrid cell identification and analysis. Tissue and PBMC segmentation were performed in QuPath using InstanSeg (version 0.1.2). For each tissue, at least three analysis regions of interest (ROIs) were drawn, spread across the specimen. For PBMCs, ROIs were drawn for secondary only control and experimental wells.

To determine pixel level thresholds for each antibody target, three spatially distinct marker negative regions were identified. For each region, the channel minimum was determined to be the level at which no signal was detected. The three channel minimums were averaged to determine the final channel minimum that would be applied for analysis. Pixel level thresholds for the blood specimen were determined using an unstained control well.

Single measurement classifiers were established for DAPI and each antigen using whole cell protein intensity averages. The minimum cell intensity average for positive signal detection of each biomarker was determined using the corresponding marker negative regions. The threshold was set as the mean cell intensity at which cells in the marker negative region were excluded from the positive detection list. For the blood specimen, classifier thresholds were set as the mean cell intensity at which cells in the unstained control well were excluded from the positive detection list. Composite classifiers including DAPI, CD45, and panCK were created to identify hybrid cells in tissue and CHCs in blood.

In PASC tumor (PASC-T), nine ROIs were analyzed for hybrid cells. After segmentation, 57,079 cells were detected. The hybrid cell composite classifier was applied, and all positive detections were manually reviewed to verify that detections contained both DAPI positive nuclei and antigen signal. Cell detections were excluded if they were incompletely captured at the ROI borders, captured fluorophore artifact that did not accurately represent protein localization, did not have positive signal for DAPI or the antigen of interest. In PASC blood (PASC-B), 49,114 cells were detected after segmentation. The hybrid cell composite classifier was applied to PASC-B, and all detections were manually reviewed with the same criteria as PASC-T.

To determine thresholds for positive signal detection for squamous and adenomatous classification and analysis in PASC-T, object classifiers for CK5 and CK8 were established based on the marker negative regions using the cytoplasmic mean intensity, aligning with expected protein localization. Composite classifiers with DAPI were made for each biomarker. In cases of positive detections in the marker negative regions when applying composite classifiers, the average of the cytoplasmic mean intensities of all positive detections was calculated and used to define the threshold for squamous versus adenomatous hybrid classification. Cell mean intensities were used for CK5 and CK8 in PASC-B squamous and adenomatous subtyping due to the loss of cell morphology during adherence of PBMCs to glass slides. Cell intensity averages were exported for every cell after all cells identified by each classifier were verified.

## Results

### PEG-PCL-NHS Synthesis and optimization of PEG chain length

We sought to directly label primary antibodies with commercially available Alexa Fluor (AF) dyes across four colors (488, 555, 647, and 750 nm) to allow for multiplexed immunostaining of multiple biomarkers. While direct antibody labeling is readily feasible, incorporation of a photocleavable linker (PCL) was necessary to facilitate signal removal for cyCIF, but introduced an additional challenge. The utilization of the hydroxyethyl photo-linker (**5**, PCL-linker, Cas: 175281-76-2) introduced the possibility of fluorescence quenching through photoinduced electron transfer (PET). To mitigate the influence of PET on fluorescence signal, we investigated the effects of polyethylene glycol (PEG) chain linker length on fluorophore photophysical properties. We hypothesized that PET quenching efficiency would decrease as the distance between the fluorophore and the PCL-linker increased. As a proof of principle, we synthesized AF488-PEG2-PCL-NHS and AF488-PEG10-PCL-NHS for primary antibody labeling (Figure 1B). Initial synthesis of AF488-PEGx-PCL-NHS (Supplementary Figure 1, Supplementary Figure 2) was completed using commercially available Boc protected PEG2 or PEG10 (**1** and **2**). The PEG2 and PEG10 were converted into symmetric PEGx anhydride intermediates **3** and **4**. Esterification of compounds **3** and **4** with the PCL-linker **5** yielded the Boc protected intermediate **6** and **7**. Acid catalyzed deprotection of **6** and **7** was followed by conjugation to AF488-NHS (**10**) to give **11** and **12**, which were treated with *N,N,N*′,*N*′-Tetramethyl-*O*-(*N*-succinimidyl)uronium tetrafluoroborate (TSTU) to yield AF488-PEG2-PCL-NHS (**13**) and AF488-PEG10-PCL-NHS (**14**).

### PEG-PCL synthetic optimization

The overall yield for **13** and **14** were <4% across the 5-step synthesis with reactions **8** and **9** being exceptionally low yielding. To improve overall synthetic yield, we generated an asymmetric acid anhydride *in situ* to decrease the required overall reaction steps to 4 and improve the esterification efficiency. As shown in Supplementary Figure 2, pivaloyl chloride (**18**) was employed to generate the asymmetric carboxylic acid anhydride intermediate, which was subsequently esterified with PCL (**5**) to yield intermediate **7**. It was observed that the *in-situ* formation of an asymmetric anhydride led to greater yields of compounds **3** and **4**. When removing the Boc protecting group, we observed that extended exposure to HCl/isopropanol yielded isopropyl ester formation at the terminal carboxylic acid. To circumvent this problem, we used 1:1 TFA (trifluoroacetic acid) and dichloromethane (DCM) conditions for Boc deprotection to obtain **8** and **9**. Overall, we were able to double the yield for the key intermediates **8** and **9**, with the yields increasing from an estimated 3.5 and 2% to 19.3 and 12.4%, respectively.

### Synthesis of AF555, AF647 and AF750 as PEG10-PCL-NHS

After synthetic approach optimization as outlined in Supplementary Figure 1, **9** was used to make AF555-(**17**), AF647-(**21**), and AF750-PEG10-PCL-NHS (**23**) constructs (Figure 1C, Supplementary Figure 3). **17** and **21** were synthesized from commercial AF555-NHS (**15**) and AF647-NHS (**19**) (Ambeed Inc, Buffalo Grove, IL and Thermo Fisher Scientific, Waltham, MA) starting materials. For, **23**, AF750 was purchased as the terminal carboxylic acid (Thermo Fisher Scientific, Waltham, MA), necessitating one additional reaction step, converting COOH to NHS ester using TSTU.

### Photophysical analysis of the effects of PEG-linker length

To evaluate the effect of PEG chain linker length on PET quenching, we measured the absorption and emission spectra, molar extinction coefficients, and quantum yields of the parent dye AF488, compound **13**, and compound **14** in PBS containing 1% DMSO. Only minimal spectral shifts were observed in both absorption and emission, indicating that the PEG–PCL modification did not introduce notable red- or blueshifts (Figure 2). As hypothesized, increasing the PEG-linker length reduced PET quenching. When the absorbance and emission of AF488, **13** and **14** were measured at equal concentrations (1 μM), compound **14** showed a higher apparent absorbance than the parent dye in our hands (Figure 2). This finding was supported by a slightly higher measured molar extinction coefficient for **14**. Notably, our measured molar absorptivity of AF488– NHS was lower than the vendor-reported value, whereas our value for **14** closely matched the vendor-reported value for AF488 (Figure 2). In contrast, both the absorbance and fluorescence intensity of **13** were substantially lower than those of AF488 and **14**. The fluorescence of **14** exhibited a moderate reduction consistent with the expected degree of PET quenching. Overall, PCL modification resulted in 58.7% and 34.8% decreases in quantum yield for PEG2 and PEG10, respectively. Based on these observations, PEG10 was selected as the optimal PEG chain length for PCL probe design and was used as the spacer for the four fluorescent probes required for PEG-PCL cyCIF imaging.

**Figure 2.**
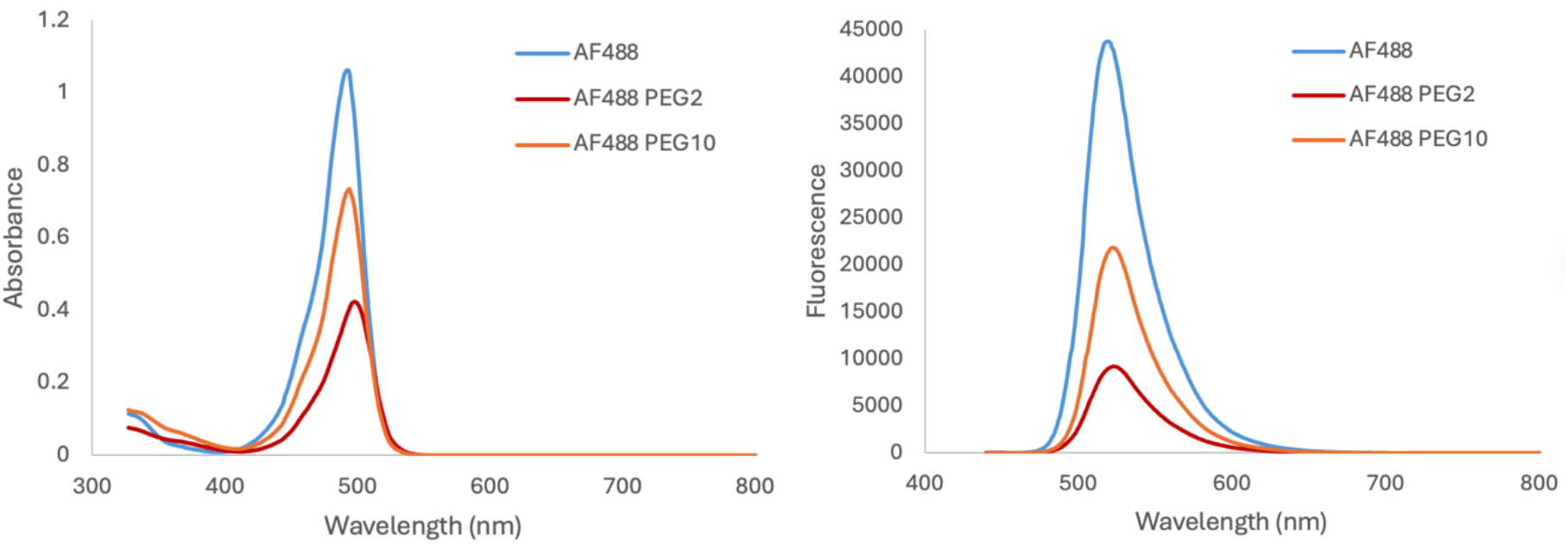
Absorbance and fluorescence spectra of parent AF488 fluorophore were compared to AF488-PEG2-PCL-NHS ester and AF488-PEG10-PCL-NHS ester to assess photoinduced electron transfer (PET) quenching. Absorbance and fluorescence of AF488-PEG10-PCL-NHS ester was closer to the parent dye demonstrating it was affected by PET quenching to a lesser extent than AF488-PEG2-PCL-NHS ester. AF488-PEG2-PCL-NHS ester and AF488-PEG10-PCL-NHS ester spectra were measured in PBS (pH 7.4) containing 1% DMSO at a concentration of 10 µM.

### Proof of concept use of PEG-PCL-FL cyCIF across primary tumor types for spatial annotation

Following optimization of PEG-linker length, we evaluated the efficacy of PEG-PCL cyCIF for annotation of cell populations and visualization of protein localization in tissues derived from distinct cancer subtypes. We selected tumors from PASC, PDAC, LUSCC, and HNSCC which exhibit a wide range of immune infiltration, squamous differentiation, and ductal structures within the tumor microenvironments (Figure 3, Supplementary Figure 4). Antibodies were selected to target epithelial cells (panCK, CK8, EpCAM, and ECAD), proteins associated with squamous differentiation (p40 and CK5), and immune populations (CD45). Additionally, we included protein targets in the epidermal growth factor (EGF) signaling pathway (EGFR, pEGFR, and pMEK) to determine if key players in cell signaling are targetable with this approach for future cell signaling analyses. Moreover, we sought to determine if PEG-PCL-FL conjugation would effectively target proteins with nuclear (p40), membranous (CD45, panCK, CK5, CK8, EpCAM, ECAD, EGFR, pEGFR), and cytoplasmic (pMEK) cellular localizations. While not every protein target was expressed in each of the four tumors, we did observe positive signal across each cellular compartment of interest (Figure 3, Supplementary Figure 4). Within the four tumor types (PASC, PDAC, LUSCC, and HNSCC) we successfully identified leukocyte infiltration, tumor epithelium, squamous cells, and kinase activation downstream of the EGF receptor. In LUSCC, HNSCC, and PASC, immune cells (Figure 3 box 1, green) were spatially close to p40+ cells (Figure 3 box 2, yellow). Additionally, these three tumors had more numerous immune clusters across the tumor compared to the PDAC tumor sample. While phosphorylated proteins, indicative of activated signaling pathways, are often challenging to target due to sample preservation, we observed robust pMEK staining across all four tumors (Figure 3 box 3, blue). This proof-of-concept application of PEG-PCL-FL conjugated antibodies established the basis for use across cancers for cyCIF analyses.

**Figure 3.**
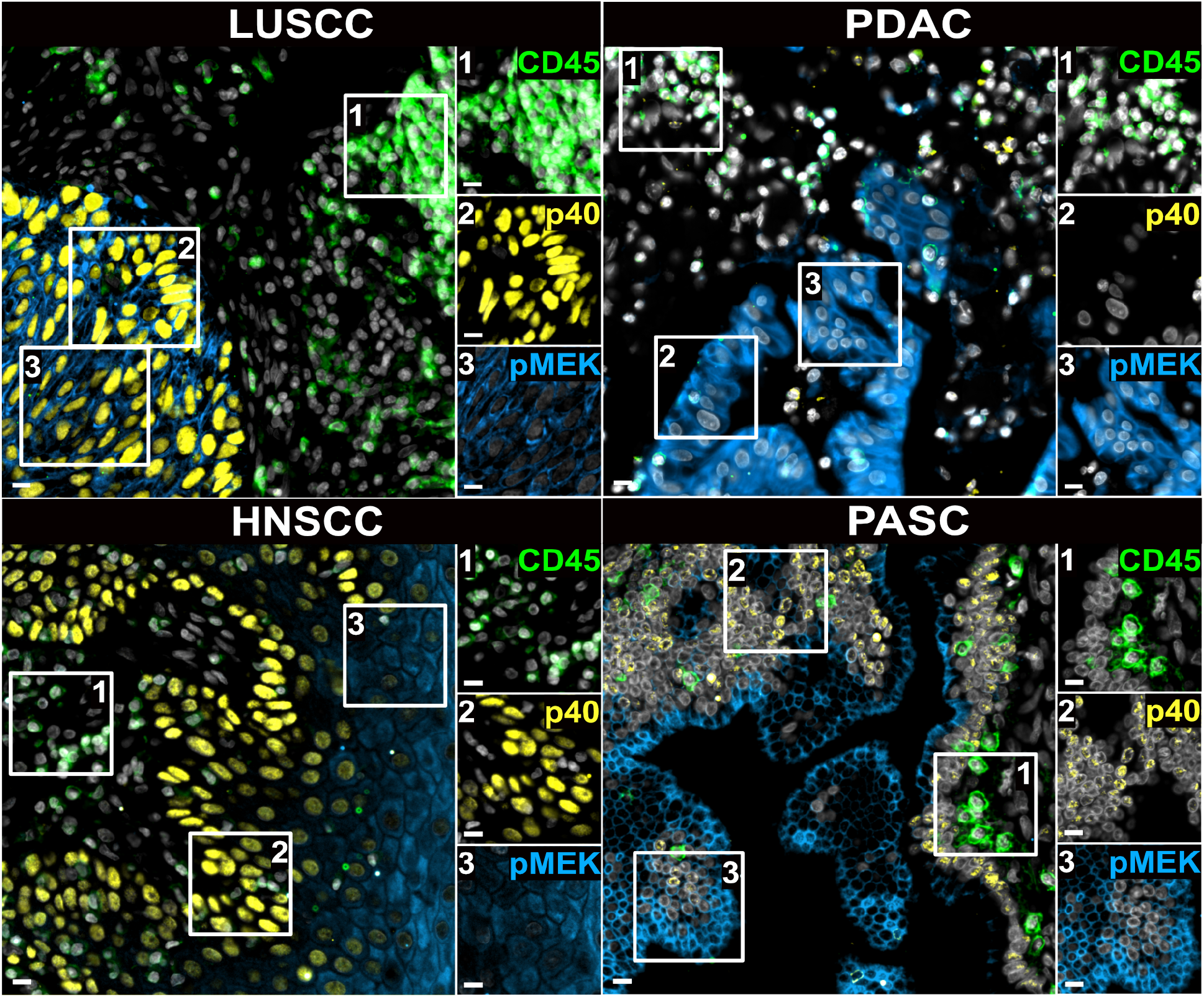
Representative images from lung squamous cell carcinoma (LUSCC), head and neck squamous cell carcinoma (HNSCC), pancreatic ductal adenocarcinoma (PDAC), and pancreatic adenosquamous carcinoma (PASC) demonstrating application of PEG-PCL-FL conjugated antibodies to membranous (CD45, green), nuclear (p40, yellow), and cytoplasmic (pMEK, blue) protein targets. White boxes featuring CD45, p40, and pMEK across the four cancer types highlight differences in cell types, immune infiltration, and cell signaling in the tumor microenvironment across cancer types. Scale bars = 100 µm.

### Proof of concept use of PEG-PCL-FL cyCIF for rare hybrid cell identification and phenotyping

PASC-T and PASC-B are patient-matched blood and tumor specimens that underwent further cyCIF analysis for hybrid cell identification and phenotyping. All tumor-based hybrid cells and blood-based circulating hybrid cells (CHCs) were identified by co-expression of CD45 and panCK in DAPI positive cells. 140 hybrid cells were identified in the PASC-T and 306 CHCs were identified in the PASC-B samples. Given the mixed squamous and adenomatous histology of PASC^27^, we sought to sub-classify identified hybrid cells as squamous or adenomatous (Figure 4). All hybrid cell expressing CK8 above the defined threshold were classified as adenomatous (Figure 4A-B). Hybrid cells that were CK8 negative, and expressed p40, CK5, or both at levels above their respective defined thresholds were classified as squamous (Figure 4C-D). Of the 140 hybrid cells identified in PASC-T, 7.7% were adenomatous and 27.9% were squamous. The remaining 64.3% were identified by panCK and CD45 co-expression, but did not express CK8, p40, or CK5 at levels above the determined thresholds for adenomatous and squamous classification. 306 CHCs were identified in PASC-B. Of these, 2.9% were adenomatous and 44.4% were squamous. Much like hybrid cells identified in the tumor, 52.6% of CHCs were identified by CD45 and panCK co-positivity without CK8, p40, or CK5 expression above the thresholds for subtype classification.

**Figure 4.**
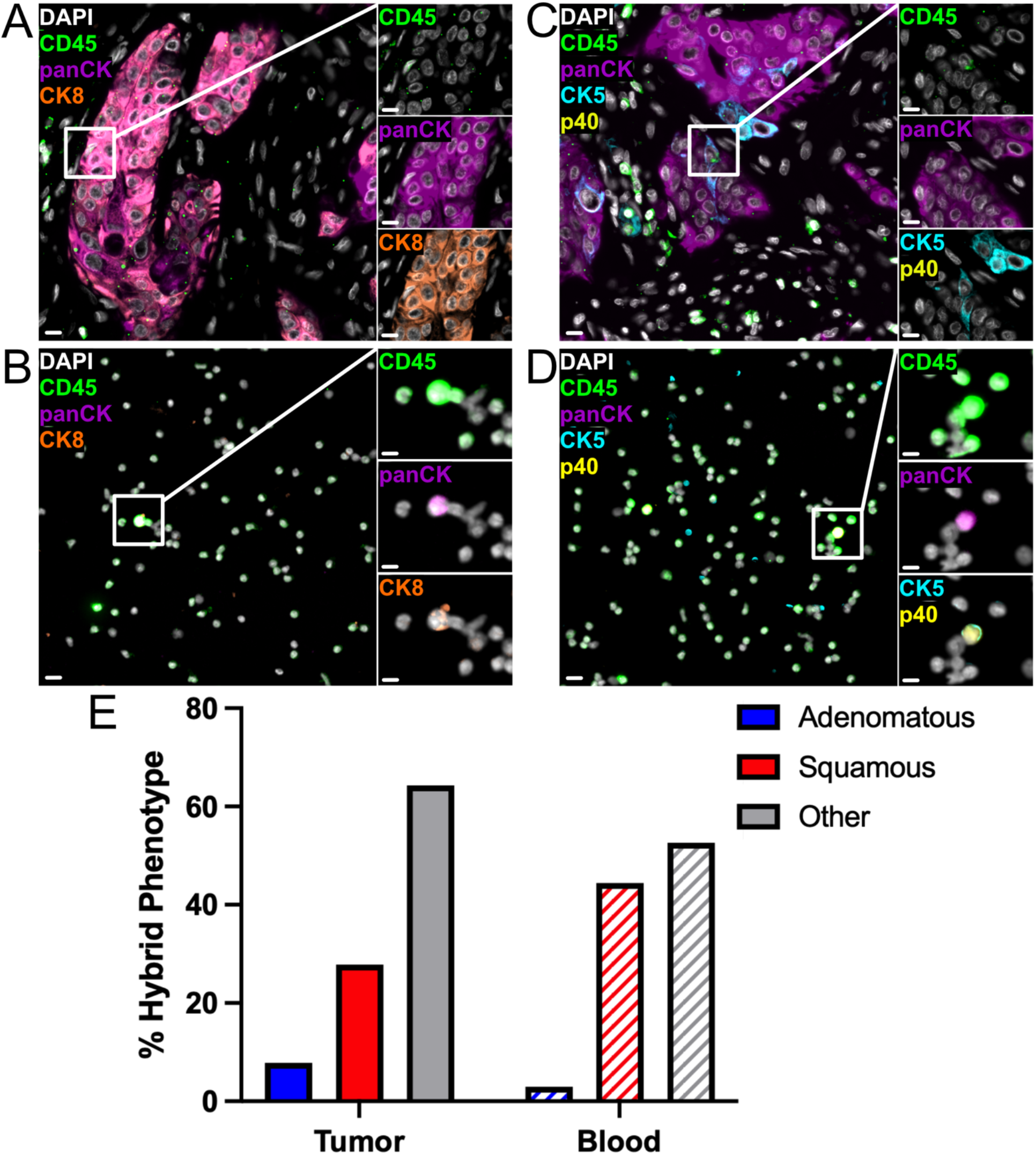
A) Adenomatous hybrid cells were identified in PASC tumor (PASC-T) by co-expression of CD45 and panCK. Adenomatous classification was determined by CK8 expression above the threshold for detection. B) Adenomatous circulating hybrid cells (CHCs) were identified in PASC blood (PASC-B) by co-expression of CD45 and panCK. Adenomatous classification in tissue hybrid cells and CHCs was determined by CK8 expression levels. C) Squamous hybrid cells were identified in PASC-T by co-expression of CD45 and panCK. D) Squamous CHCs were identified in PASC-B by co-expression of CD45 and panCK. Squamous classification was determined by CK5 and/or p40 expression levels. Scale bars on all tissue merged and individual protein images represents 100 µm. Scale bars on all blood merged images represent 100 µm and 50 µm on individual protein images. E) In PASC-T and PASC-B, squamous hybrids comprise a higher percentage than adenomatous hybrids of all identified hybrids. The relative similarity of cell type classification between primary tumor and blood specimens demonstrates that blood samples are representative metric for hybrid cell phenotyping of primary tumors.

## Discussion

Multiplexed immunofluorescence tools are a powerful method for elucidating spatial proteomics and complex tumor biology. However, these platforms routinely face challenges in maintaining specimen integrity due to the harsh methods required for signal removal between rounds of staining and imaging. In this study, we validated and demonstrated that PCLs can be implemented for gentle signal removal as part of a fluorophore conjugate to directly label primary antibodies with fluorophores, where PEG spacers were utilized to facilitate maintain emitted fluorescence signal. Use of PEG-PCL-FL conjugated antibodies permitted UV cleavage of fluorescent signal between rounds of staining and imaging, minimizing sample degradation across staining rounds that is common to other cyCIF imaging methodologies. Gentle fluorophore cleavage is a high priority when performing cyCIF on PBMC specimens because adhered PBMCs do not benefit from an extracellular matrix, making them more susceptible to sample degradation with standard fluorophore bleaching or antibody dissociation techniques for cyCIF signal removal. Additionally, we anticipate application of PEG-PCL-FL antibody conjugates to cyCIF will maintain the ability to quantify protein expression for cell type classification that is inherent to the utility of direct immunofluorescence. Implementation of the PEG10 spacer ensured maintenance of AF dye photophysical properties (Figure 2), preserving absorbance and emission wavelengths as well as high extinction coefficient and quantum yield permitting high signal to background ratio (SBR) imaging. High SBR imaging is especially important in the 488 and 555 nm imaging channels, which are affected by high autofluorescence signal in tissue and PBMC samples. Methods to minimize autofluorescence signal in tissue and PBMC samples is an area of active investigation. Decreasing autofluorescence would further enhance cyCIF SBR, improving ease and quality of subsequent analyses.

We successfully applied cyCIF with PEG-PCL-FL conjugated antibodies to varied cancer types including PASC, PDAC, LUSCC, and HNSCC, probing nuclear, cytoplasmic, and membrane protein targets (Figure 3, Supplementary Figure 4). Our antibody panel encompassing CD45, a pan-leukocyte antigen, numerous epithelial proteins (EpCAM, ECAD, panCK, CK5, and CK8) and cell signaling pathway proteins (EGFR, pEGFR, and pMEK) enabled us to identify cell types within the tumor microenvironment, tumor architectural structures and cell phenotypes across cancer subtypes. PEG-PCL-FL conjugated antibodies represent an optimal approach to identify and subtype rare hybrid cells in PASC tumor and blood.

We demonstrated that our optimized cyCIF and analysis pipeline can accurately identify hybrid cells in tumor and blood specimens by co-expression of CD45 and panCK. Through our analysis of PASC-T and PASC-B, we further sub-classified hybrid cells as squamous or adenomatous in origin by their expression of CK8, p40, and CK5. The relative percentages of squamous and adenomatous hybrid cells identified were similar in PASC-T and PASC-B indicating that CHCs are a promising noninvasive liquid biopsy that can be analyzed from a blood draw to provide insight into the composition and phenotype of the primary tumor. This result supports development of expanded cyCIF panels enabling further hybrid cell phenotyping to define the underlying tumor biology of aggressive cancers such as PASC, with potential to inform treatment decisions and stratify patients for new and existing clinical trials.

One limitation of our technology is the variable success of NHS ester antibody conjugation as not all antibody clones yield functional conjugates. Another limitation is the potential presence of free, unconjugated PEG-PCL-FL with the PEG-PCL-FL conjugated antibodies. While antibodies were purified to minimize unconjugated PEG-PCL-FL, some aggregates remained, requiring thorough image review to remove cells visibly containing fluorophore aggregates after pixel level histogram thresholds and classifier thresholds were applied. Future exploration of PEG-PCL-FL conjugation with site-click chemistry could minimize this challenge due to increased antibody conjugation efficiency. One additional limitation of this work are limitations in cell segmentation, resulting in over-segmenting or under-segmenting cells in tumor and blood specimens. Use of DAPI staining as well as cytoplasmic and membrane proteins enhances segmentation quality by better defining cell boundaries compared to DAPI alone but does not fully prevent over- or under-segmentation. In PBMC specimens, segmentation quality can be affected by the homogeneity of PBMC spread across each well on the glass slide.

## Conclusion

Significant advances have been made in molecular immunofluorescence imaging techniques, yet technical limitations remain hindering broad application of cyCIF on diverse biologic specimens. cyCIF emerged as a tool applicable across disease pathologies for generating highly multiplexed images through multiple rounds of antibody staining, imaging, and signal removal. However, current methods for fluorophore removal impose harsh conditions, altering antigenicity and tissue integrity, thus limiting utility on fragile biologic specimens. Our previously validated Ab-oligo cyCIF methodology addressed these concerns by adding a photocleavable linker for fluorophore removal using UV light treatment, enabling us to successfully implement cyCIF on tumor and peripheral blood specimens. However, this method remained limited for rare cell detection by nonspecific nuclear accumulation that resulted in false positive detection of nuclear targets.

To address the limitations of Ab-oligo cyCIF, we developed an optimal PEG-PCL-FL conjugation strategy to directly label primary antibodies. We tested varied length PEG spacers (PEG2 and PEG10) to determine which spacer length would minimize PCL fluorophore quenching due to PET and yield fluorescence intensities similar to that of the parent fluorophore. We determined that PEG10 spacers conjugated between the PCL and fluorophore most effectively maintained fluorophore brightness and applied this to antibody conjugation with spectrally distinct Alexa Fluor dyes (AF488, AF555, AF647, AF750). Using our cyCIF with PEG-PCL-FL directly labeled antibodies, we constructed images of the primary tumor across four cancer types including PASC, PDAC, LUSCC, and HNSCC. We demonstrated that direct antibody labeling portrays the spatial localization of target proteins in each cellular compartment (membrane, cytoplasm, and nucleus) and enabled use of protein expression for subtype classification on a single-cell basis. We further utilized PEG-PCL-FL cyCIF to identify, phenotype, and classify rare cell populations, including neoplastic-immune hybrid cells, which are present in both tumor and peripheral blood. Here we demonstrate that PEG-PCL cyCIF is an enhanced and robust flexible cyCIF platform with significant promise to advance phenotypic analyses of tumor tissue and PBMC specimens across cancers to inform critical clinical decisions.

## Acknowledgements

We would like to thank the Advanced Light Microscopy Core at Oregon Health and Science University (OHSU), the OHSU Knight Biolibrary, and the Brenden-Colson Center for Pancreatic Care. This research utilized OHSU’s Advanced Computing Center Services computational infrastructure supported by the Office of Research Infrastructure Programs, Office of the Director of the National Institutes of Health under Award Number S10OD034224. Figures were created using BioRender, Affinity Design 2, and ChemDraw.

## Author Contributions

Conceptualization, A.Z., C.N., F.B., M.H.W., L.G.W., S.L.G.

Methodology, A.Z., C.N., F.B., G.S.M., J.J., T.J.E., C.R., D.R., M.H.W., L.G.W., S.L.G.

Formal Analysis, A.Z., C.N., F.B., D.R.

Data Curation, A.Z., C.N., F.B.

Writing – original draft preparation, A.Z., C.N., F.B., J.J., M.H.W., L.G.W., S.L.G.

Writing – reviewing and editing, all authors.

Resources, S.M.G., A.K., M.H.W., L.G.W., S.L.G.

## References

1. R. L. Siegel et al., “Cancer statistics, 2026,” CA: A Cancer Journal for Clinicians 76(1)(2026). 10.3322/caac.70043.

2. American Cancer Society, “Cancer Facts & Figures 2026,” (2026).

3. B. Seliger and C. Massa, “CyTOF as a suitable tool for stratification and monitoring of cancer patients,” Journal of Translational Medicine 23(1)(2025). 10.1186/s12967-025-06764-0.

4. J. Ptacek et al., “Multiplexed ion beam imaging (MIBI) for characterization of the tumor microenvironment across tumor types,” Laboratory Investigation 100(8), 1111–1123 (2020). 10.1038/s41374-020-0417-4.

5. M. Angelo et al., “Multiplexed ion beam imaging of human breast tumors,” Nature Medicine 20(4), 436–442 (2014). 10.1038/nm.3488.

6. Y. Zhou et al., “Metal-detection based techniques and their applications in metallobiology,” Chemical Science 15(27), 10264–10280 (2024). 10.1039/d4sc00108g.

7. C. Giesen et al., “Highly multiplexed imaging of tumor tissues with subcellular resolution by mass cytometry,” Nat Methods 11(4), 417–422 (2014). 10.1038/nmeth.2869.

8. R. M. Levenson, A. D. Borowsky and M. Angelo, “Immunohistochemistry and mass spectrometry for highly multiplexed cellular molecular imaging,” Laboratory Investigation 95(4), 397–405 (2015). 10.1038/labinvest.2015.2.

9. E. R. Parra, A. Francisco-Cruz and I. I. Wistuba, “State-of-the-Art of Profiling Immune Contexture in the Era of Multiplexed Staining and Digital Analysis to Study Paraffin Tumor Tissues,” Cancers 11(2), 247 (2019). 10.3390/cancers11020247.

10. J. R. Lin et al., “Highly multiplexed immunofluorescence imaging of human tissues and tumors using t-CyCIF and conventional optical microscopes,” Elife 7](2018). 10.7554/eLife.31657.

11. M. J. Gerdes et al., “Highly multiplexed single-cell analysis of formalin-fixed, paraffin-embedded cancer tissue,” Proceedings of the National Academy of Sciences 110(29), 11982–11987 (2013). 10.1073/pnas.1300136110.

12. E. C. Stack et al., “Multiplexed immunohistochemistry, imaging, and quantitation: a review, with an assessment of Tyramide signal amplification, multispectral imaging and multiplex analysis,” Methods 70(1), 46–58 (2014). 10.1016/j.ymeth.2014.08.016.

13. J.-R. Lin, M. Fallahi-Sichani and P. K. Sorger, “Highly multiplexed imaging of single cells using a high-throughput cyclic immunofluorescence method,” Nature Communications 6(1), 8390 (2015). 10.1038/ncomms9390.

14. P. Zrazhevskiy, L. D. True and X. Gao, “Multicolor multicycle molecular profiling with quantum dots for single-cell analysis,” Nature Protocols 8(10), 1852–1869 (2013). 10.1038/nprot.2013.112.

15. P. Zrazhevskiy and X. Gao, “Quantum dot imaging platform for single-cell molecular profiling,” Nature Communications 4(1), 1619 (2013). 10.1038/ncomms2635.

16. J. R. Lin et al., “Cyclic Immunofluorescence (CycIF), A Highly Multiplexed Method for Single‐cell Imaging,” Current Protocols in Chemical Biology 8(4), 251–264 (2016). 10.1002/cpch.14.

17. G. Glass, J. A. Papin and J. W. Mandell, “Simple: A Sequential Immunoperoxidase Labeling and Erasing Method,” Journal of Histochemistry & Cytochemistry 57(10), 899–905 (2009). 10.1369/jhc.2009.953612.

18. N. P. McMahon et al., “Flexible Cyclic Immunofluorescence (cyCIF) Using Oligonucleotide Barcoded Antibodies,” Cancers 15(3), 827 (2023). 10.3390/cancers15030827.

19. J. A. Jones et al., “Oligonucleotide conjugated antibody strategies for cyclic immunostaining,” Scientific Reports 11(1)(2021). 10.1038/s41598-021-03135-9.

20. N. P. McMahon et al., “Oligonucleotide conjugated antibodies permit highly multiplexed immunofluorescence for future use in clinical histopathology,” Journal of Biomedical Optics 25(05), 1 (2020). 10.1117/1.jbo.25.5.056004.

21. M. E. McCarthy et al., “Increasing Signal Intensity of Fluorescent Oligo-Labeled Antibodies to Enable Combination Multiplexing,” Bioconjug Chem 35(7), 1053–1063 (2024). 10.1021/acs.bioconjchem.4c00246.

22. C. E. Gast et al., “Cell fusion potentiates tumor heterogeneity and reveals circulating hybrid cells that correlate with stage and survival,” Science Advances 4(9), eaat7828 (2018). 10.1126/sciadv.aat7828.

23. M. S. Dietz et al., “Relevance of circulating hybrid cells as a non-invasive biomarker for myriad solid tumors,” Scientific Reports 11(1)(2021). 10.1038/s41598-021-93053-7.

24. C. P. Corkum et al., “Immune cell subsets and their gene expression profiles from human PBMC isolated by Vacutainer Cell Preparation Tube (CPT™) and standard density gradient,” BMC Immunology 16(1)(2015). 10.1186/s12865-015-0113-0.

25. I. Johnson and M. Spence, “The Molecular Probes Handbook: A Guide to Fluorescent Probes and Labeling Technologies,” Life Technologies, [https://www.thermofisher.com/us/en/home/references/molecular-probes-the-handbook/tables/fluorescence-quantum-yields-and-lifetimes-for-alexa-fluor-dyes.html]

26. J. L. Muhlich et al., “Stitching and registering highly multiplexed whole-slide images of tissues and tumors using ASHLAR,” Bioinformatics 38(19), 4613–4621 (2022). 10.1093/bioinformatics/btac544.

27. Y.-F. Feng et al., “110 Patients with adenosquamous carcinomas of the pancreas (PASC): imaging differentiation of small (≤ 3 cm) versus large (> 3 cm) tumors,” Abdominal Radiology 44(7), 2466–2473 (2019). 10.1007/s00261-019-01989-2.

