## Supplemental Data for "Cyclic immunofluorescence platform using photocleavable linkers for direct antibody labeling enables cancer phenotyping"

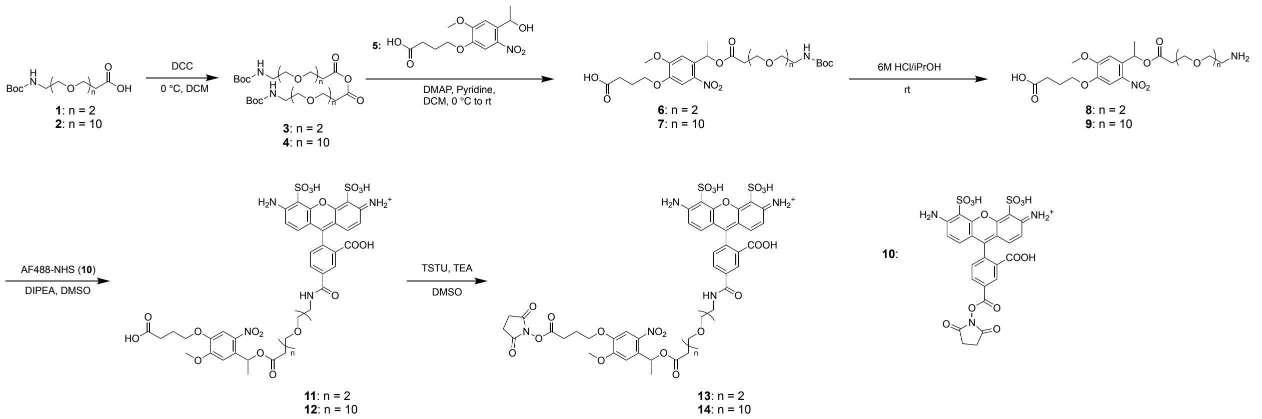


**Supplementary Figure 1.** General synthetic route to AF488-PEGx-PCL-NHS ester. PEGx-PCL compounds (**8**, **9**) were synthesized using commercially available PEG chains (**1**, **2**) and PCL (**5**). The deprotected compounds (**8**, **9**) were conjugated to AF488-NHS (**10**) yielding **11**, and **12** bearing terminal carboxylic acid. TSTU was used to furnish final products **14** and **15**. (PCL, 4-(4-(1-Hydroxyethyl)-2-methoxy-5-nitrophenoxy)butanoic acid (compound **5**); DCC, *N*,*N*'-Dicyclohexylcarbodiimide; DCM, dichloromethane; DMAP, 4-Dimethylaminopyridine; TEA, triethylamine; TSTU, *N*,*N*,*N*',*N*'-Tetramethyl-*O*-(*N*-succinimidyl)uronium tetrafluoroborate; DMSO, dimethyl sulfoxide; Boc, tert-butyloxycarbonyl.)

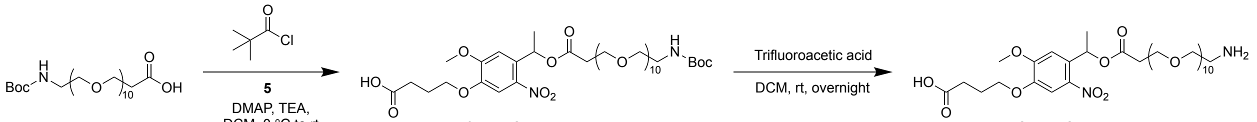


**6**: n = 2
**7**: n = 10

**8**: n = 2
**9**: n = 10

**Supplementary Figure 2.** Synthetic route to PEG10-PCL (**9**) using a mixture of pivalolyl chloride (**18**), t-Boc-N-amido-PEG10-amine (**2**), and PCL (**5**). TEA, triethylamine; DMAP, 4-dimethylaminopyridine; DCM: dichloromethane.

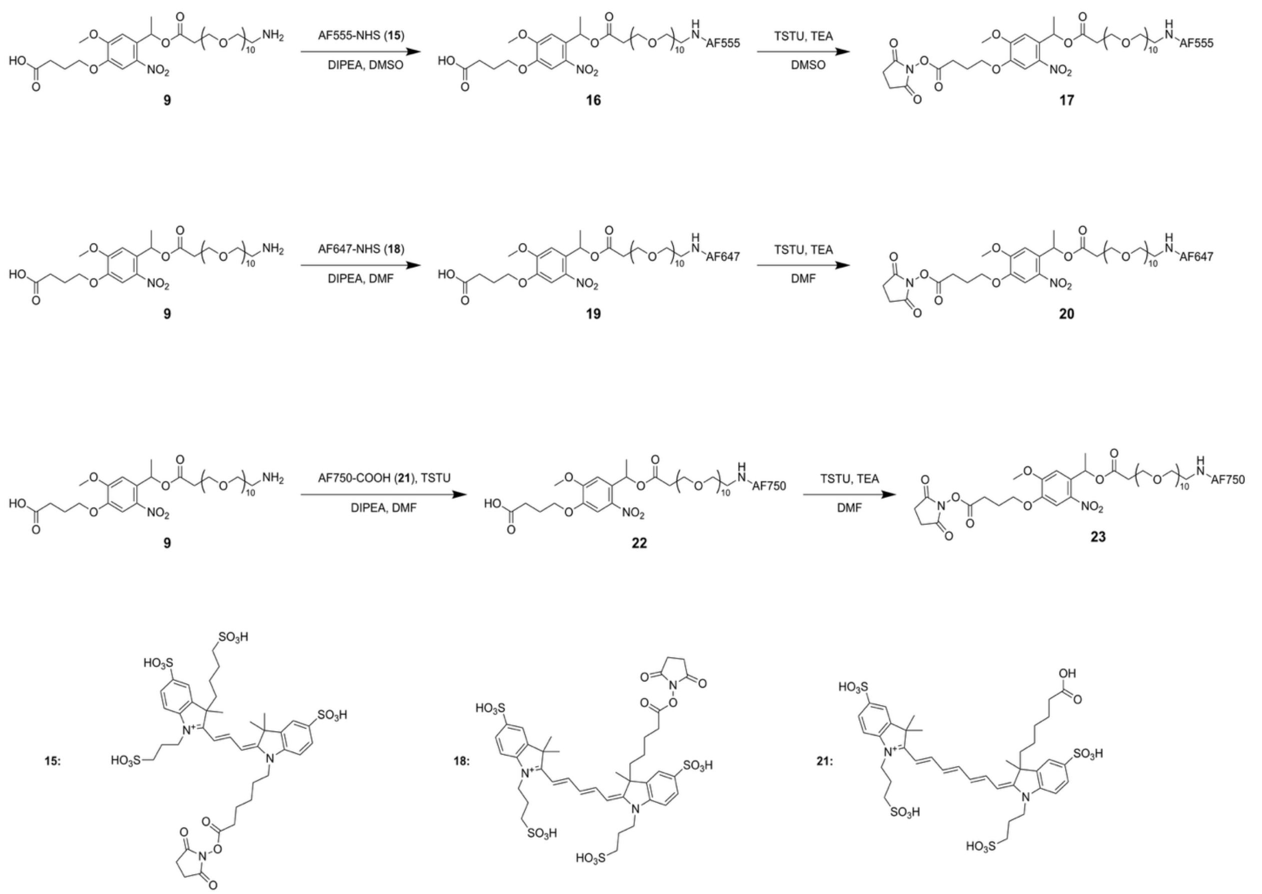


**Supplementary Figure 3**. Synthetic routes to PEG10-PCL NHS esters for AF555, AF647, and AF750. Using PEG10-PCL (**9**) synthesized following the protocol outlined in scheme 2 compounds **17**, **20**, and **23** were synthesized from commercial AF dye intermediates over two steps. TEA, triethylamine; TSTU, N,N,N',N'-Tetramethyl-O-(N-succinimidyl)uronium tetrafluoroborate; DMF, dimethyl formamide; DMSO, dimethyl sulfoxide.

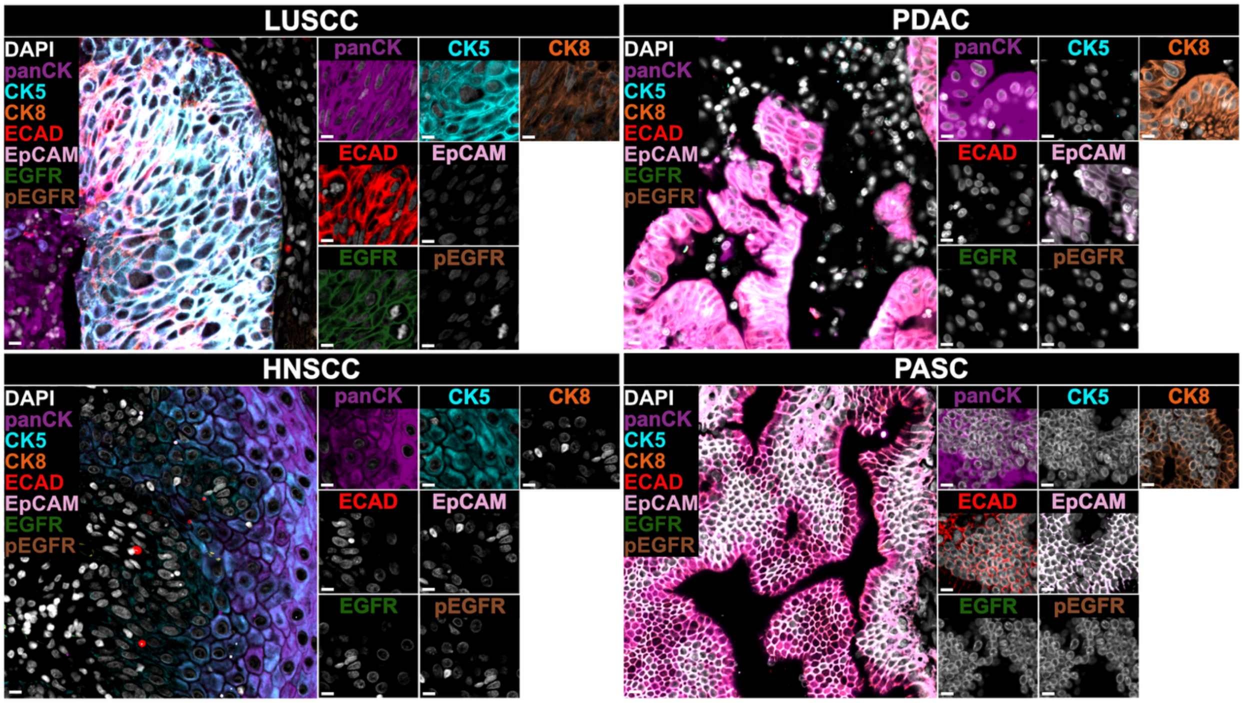


**Supplementary Figure 4.** Representative images from lung squamous cell carcinoma (LUSCC), head and neck squamous cell carcinoma (HNSCC), pancreatic ductal adenocarcinoma (PDAC), and pancreatic adenosquamous carcinoma (PASC) demonstrating additional membranous and cytoplasmic proteins targeted with PEG-PCL-FL conjugated antibodies. Scale bars = 100 µm.

**Supplementary Table 1.** Photophysical data of AF-labeled fluorescence probes and AF dyes.

| Compound[a] | $\lambda_{abs}$ (nm) | $\lambda_{em}$ (nm) | $\varepsilon^{[b]}$ ($10^4$ $M^{-1}$ $cm^{-1}$) | $\Phi_f^{[c]}$ |
|---|---|---|---|---|
| AF488[d] | 492 | 519 | 6.3 | 0.92 |
| AF488-PEG2-PCL-NHS | 498 | 523 | 4.7 | 0.38 |
| AF488-PEG10-PCL-NHS | 494 | 523 | 5.1 | 0.60 |
| AF555[e] | 550 | 569 | 14.6 | 0.10 |
| AF555-PEG10-PCL-NHS | 550 | 569 | 8.8 | 0.11 |
| AF647[f] | 647 | 670 | 27.3 | 0.33 |
| AF647-PEG10-PCL-NHS | 649 | 673 | 34.6 | 0.21 |
| AF750[g] | 749 | 775 | 29 | 0.12 |
| AF750-PEG10-PCL-NHS | 751 | 776 | 6.5 | 0.17 |

[a] All compounds were dissolved in PBS (pH 7.4) containing 1% DMSO (10 μM). [b] ε: extinction coefficients were calculated using Beer's Law by plotting absorbance values against 5 different concentrations to obtain the slope. [c] Fluorescence quantum yields. [d], [e], and [f] Quantum yields of AF488, AF555, and AF647 were obtained from ThermoFisher Scientific. [g] Extinction coefficient and quantum yield of AF750 were obtained from ThermoFisher Scientific.
